# Langevin dynamics simulation of protein dynamics in nanopores at microsecond timescales

**DOI:** 10.1101/2021.06.21.449278

**Authors:** J. P. Mahalik, Jeffrey Cifello, Murugappan Muthukumar

## Abstract

With rapid advancement in the fields of nanopore analysis of protein, it has become imperative to develop modeling framework for understanding the protein dynamics in nanopores. Such modeling framework should include the effects of electro-osmosis, as it plays significant role during protein translocation in confinement. Currently, the molecular dynamics simulations that include the hydrodynamic effects are limited to a timescale of few 100 ns. These simulations give insight about important events like protein unfolding which occurs in this timescale. But many electrophoresis experiments are limited by a detector resolution of ~ 2.5 *μs*. Analytical theory has been used to interpret protein dynamics at such large timescale. There is a need for molecular modeling of more complex environment and protein shapes which cannot be accounted for by analytical theory. We have developed a framework to study globular protein dynamics in nanopores by using langevin dynamics on a rigid body model of protein and the hydrodynamics is accounted by analytical theory for simple cylindrical nanopore geometry. This framework has been applied to study the dynamics of Ubiquitin translocation in SiN_*x*_ nanopore by Nir et al^26^. They have reported 7 times decrease in average dwell time of the protein inside the nanopore in response to a small change in pH from 7.0 to 7.2 and the modification of protein charge was attributed for such drastic change. Closer examination using our simulation revealed that the electro-osmotic effects originating due to very small change in the surface electrostatic potential of the nanopore could lead to such a drastic change in protein dynamics.

## I. INTRODUCTION

Nanopore technology provides exciting opportunities for characterizing proteins, it probes them at a single molecule level, typically requires very small amount of sample (~ nM), and is a label free method^1,2^. This technology has been used in many applications, such as in protein sensing^2,3^, monitoring conformational changes of proteins such as folding/unfolding^4–8^, and for self-assembly of proteins inside a nanopore^9–11^. Recently, there has been a lot of development and numerous opportunities in nanopore analysis of proteins, as evident from the large number of review articles in this particular field in just the past few years^1–3,12–16^. Protein analysis using nanopores is an indirect method wherein the structural and/or dynamic properties of a protein molecule are interpreted through ionic current trace. When an electric field is applied across a nanopore immersed in a hypertonic solution, a steady baseline open pore ionic current (~ pA to nA) is observed. When a protein of size at least comparable to that of the nanopore diameter moves through it, total ionic flux is reduced, leading to a detectable reduction in the ionic current for certain duration of time. This current is referred as blockade current and it has two important characteristics: blockade current depth with respect to the open pore current, and the blockade duration (there could be multiple blockade depth levels). Many factors could influence the conformation and motion of a protein molecule inside the nanopore, such as interaction of protein molecule with the nanopore, confinement effect, and flow field of the aqueous media. Interpreting how protein structure and dynamics may be influenced by so many factors can be challenging based on just two sets of information: blockade current depth and blockade duration. A complimentary direct observation technique may facilitate such interpretation. All atomistic Molecular dynamics (MD) has been used to understand conformational changes of protein inside nanopores^17–19^, translocation dynamics^20–22^, protein sensing^23^, which typically happen in a timescale of less than or of the order of 100 ns. Coarse grain models have also been used to investigate protein translocation. Using Martini CG forcefield Haridasan et al^24^ investigated the translation of streptavidin though nanopores of different diameter, demonstrating how strong confinement can slow down protein dynamics due to strong protein-pore interactions. This simulation was of the timescale of few 100 ns. However, electrophoresis experiments are limited by their detector resolution at ~ 2.5 *μs*^25,26^, the lowest dwell time reported for RNase^25^ and Ubiquitin^26^ is ~ *μs.* Besides the average dwell time, the dwell time distribution spans tens of *μs*, if not hundreds of *μs*. Analytical theory has been used to interpret the protein dynamics at such large timescales^25,27^. However, analytical calculations may not be applied to all situations, especially for more complex environment and protein shapes, where the details of the system such as shape of the protein can significantly impact the dynamics. A generalized framework is needed for probing protein dynamics in confinement at a timescale of *μs* or higher. Ustach and Faller^28^ proposed raspberry model for protein-like particle dynamics in cylindrical confinement at large timescales. They calculated the hydrodynamic resistances of ellipsoid raspberries to translational and rotational motion using lattice Boltzmann and highlighted the need for considering more complex rigid shapes.

A protein molecule experiences different kinds of forces during its expedition through a nanopore. Unlike DNA or RNA, a protein molecule is typically made up of a mixture of positive and negative charged residues. The externally applied electric field exerts electrophoretic force on positively charged residues in one direction and pushes the negatively ones in the opposite direction, resulting in a net translational and rotational motion of the protein in a particular direction. Secondly, depending on the nature of charge of the nanopore and the polarity of the applied voltage, electroosmotic driving force may become significant, leading to either enhancement or reduction of the protein velocity. Also, the interaction of the protein with the nanopore walls and the confinement effects of the nanopore may influence the motion of the protein molecule. In addition to influencing the dynamics of the protein, the confinement effect of the nanopore may cause structural changes in the protein molecule such as unfolding^17–19^. We will consider only the simplest case when the protein moves through the nanopore without any significant conformational changes^2^. Muthukumar^27^ modeled the effects of counterion cloud, external electric field, confinement effect inside nanopore and protein-pore interaction on the dynamics of protein translocation through nanopore using analytical theory. Without adequately accounting for all this factors, the interpretation from the single molecule electrophoresis experiments can be misleading. For example, at high salt the effective charge on a protein molecule (*Q*) could be reduced by about an order of magnitude and the diffusivity of the protein (*D*) could be reduced significantly due to the confinement effects and the protein-pore attraction. Simply using the Einstein equation (*μ* = *QD/k_B_T*) for interpreting the protein dynamics can be erroneous. Where *μ* is the electrophoretic mobility of the protein, *k_B_T* is the Boltzmann constant times the absolute temperature, *Q* is typically taken to be overall charge of the protein without accounting for its counterion cloud, and *D* is generally approximated from Stokes-Einstein relation for a sphere in a free medium. This analytical approach provides a good starting point for estimating the effects of some of the factors on the dynamics of protein during translocation through a nanopore. Specific details can become important in many situations, for example, Liu et al^29^ investigated the effect of wettability condition inside the nanopore on the CG protein translocation velocity, stronger attraction between the water molecules and the nanopore can slow down the protein dynamics significantly in hydrophilic nanopores instead of protein-nanopore interactions. Such insight could be obtained by molecular modeling, otherwise it could have been wrongly attributed to some other factor.

Nir et al^26^ investigated translocation of Ubiquitin through a SiN_*x*_ nanopore of diameter comparable to its size (3.6 nm diameter nanopore). They observed that the average protein dwell time decreased by about a factor of 7, with a small change in pH from 7.0 to 7.2 from 18.4 *μs* to 2.6 *μs*, respectively. For pH values higher than 7.2, the translocation events were too fast to be detected. For pH values lower than 7.0, the capture rate dramatically dropped, making it difficult for statistical analysis. Such a drastic response of translocation time to such a small change in pH was attributed to the modification of the protein charge in response to change in pH. However, analysis of the Ubiquitin sequence^30^ revealed that it has only one Histidine residue and that is also expected to be in deprotonated state for pH range 7.0-7.2^31^. Therefore, the electrostatic effects must be originating from the nanopore. At a high salt concentration of 1.5 M KCl, we expect that the electrostatic interactions will be highly screened. Most likely the drastic change in protein dynamics is originating from electroosmotic interactions. In order to account for hydrodynamics in such systems, generally explicit solvent molecules are considered atomistically^20–22^ or CG^24,29^, or less often through hybrid model: coupling MD of the biomolecule with hydrodynamic interactions from Lattice-Boltzmann (LB) calculations^32^. Such hybrid model can still be computationally expensive, requiring numerical LB calculations over the grids at different timesteps. For simplified geometries such as a cylindrical nanopores of sufficiently large diameter, continuum approximations can be used to obtain fluid velocity profile as a function of nanopore surface potential, applied electric field, fluid viscosity, nanopore diameter and salt concentration^33^. At sufficiently low Reynolds number, we can assume that the biomolecule experiences the electro-osmosis force but it does not influences the fluid flow profile around it. Such simplified assumption leads to only one calculation of fluid flow profile in the beginning of MD calculations, the fluid velocity profile can then be related to particle dynamics through langevin dynamics. Another computational challenge is accounting for different kinds of interactions (bonds, angular, dihedral, etc) between the residues during translocation. If the protein unit is small enough and globular to slide through the nanopore and the external forces are weak enough such that the internal fluctuations within the protein structure can be neglected, then we can model the protein unit as a rigid body of any arbitrary shape (depending on the Protein Data Bank (PDB) structure) instead of restricting to symmetric shapes^28^. The residues can interact with the nanopore through short-range and long-range pairwise interactions based on the nature of the residues.

We used langevin dynamics simulations to study the dynamics of electrophoretically driven Ubiquitin through a SiN_*x*_ nanopore of diameter 3.6 nm^26^. The electro-osmotic effects were accounted for analytically and the protein is modeled as a rigid body. With these assumptions, we were able to monitor Ubiquitin dynamics inside the nanopore upto tens of *μs* and we demonstrate that the electro-osmosis can drastically reduce the dwell time of the Ubiquitin. A surface potential of ~ *mV* can have a significant impact on the dynamics, changing the qualitative dynamics of Ubiquitin from diffusion like to drift like.

## II. MODEL AND SIMULATION DETAILS

Simulations were performed at two different length/time scales: all atomistic molecular dynamics (MD) simulations and coarse grained langevin dynamics (LD) simulations. All atomistic MD simulation was performed on a Ubiquitin molecule immersed in bulk 1.5 M KCl aqueous solution in nanosecond timescale. The stability of the protein structure was determined in the bulk without and with externally applied electric field. The root mean square deviation (RMSD) of the protein backbone as a function of time was used to gauge the stability of the protein in the timescale of nanoseconds. MD simulations were performed with CHARMM protein potential and TIP3P water model using NAMD^34^ software. Since the protein was found to be stable in the bulk (both without and with the externally applied electric field), the next stage of the simulation was conducted at a timescale of microseconds on a coarse grained (CG) model of the protein, assuming a rigid protein structure. The coarse graining was done at the residue level, each CG bead represents one residue. A rigid protein structure implies that the relative position of every residue with respect to each other is kept fixed during the motion of the protein unit. Such a simplification allows turning off of interaction among beads within the protein, their dynamics is determined through their interaction with the environment. The second part of the simulation is described in detail below:

Langevin dynamics simulation was used to investigate the Ubiquitin dynamics in the nanopore, using LAMMPS^35^. The Langevin equation for the *j^th^* component of the position vector of the *i^th^* bead (*r_ij_*) is given by,

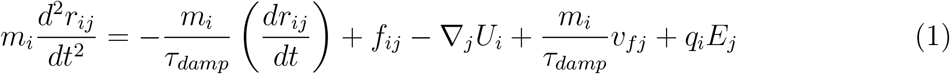

where *m_i_* is the mass, *τ_damp_* is the damping factor, and *v_fj_* is the *j^th^* component of the local fluid velocity acting on the *i^th^* bead. *f_ij_* is the *j^th^* component of the random force acting on the *i^th^* bead obeying the fluctuation-dissipation theorem with its magnitude proportional to 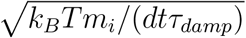 (*k_B_T* being the Boltzmann constant times the absolute temperature and *dt* is the timestep), its magnitude and direction being randomized using random numbers. *m_i_* is taken to be same for all the beads and Stokes’ law was used to estimate the *τ_damp_*, *m_i_*/*τ_damp_* = 3*πησ*, where *η* is the viscosity of the water at 298 K equal to 1.0 × 10^-3^ poise. *τ_damp_* is estimated to be 51 femtoseconds, based on these parameters. Refer Table I for *m_i_* and *σ* (diameter of a CG residue) values. The second last term on the RHS of Eq. 1 is the electro-osmotic force due to fluid velocity *v_fj_* and the last term is the electrophoretic force experienced by a bead possessing charge *q_i_* due to external electric field *E_j_*.

**TABLE I:** Fundamental parameters in the simulation from real to LJ units

| Parameter | Real Units | LJ units |
| --- | --- | --- |
| Mass ($m_0$) | 110 g/mol (Average mol wt of residue) | 1 |
| Length ($\sigma$) | 3.8 Å (Diameter of CG residue) | 1 |
| Time ($\tau$) | 2.53 ps | 1 |
| Energy ($\epsilon$) | 1 $k_B T$ at 298 K | 1 |
| Force ($F$) | 10.82 pN | 1 |
| Charge ($q$) | 1 electronic charge | 12.14 |
| Electric field ($E$) | 1 V/Å | 0.0821 |

*U_i_* is the net potential on the *i^th^* bead due to contributions from the repulsive excluded volume interactions and the screened Coulombic interactions. The excluded volume interaction is modeled as Lennard Jones (LJ) potential

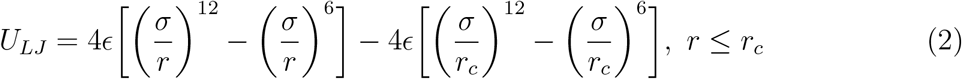

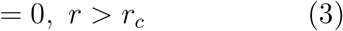

where *σ* and *ϵ* are parameters. We have chosen the cut-off *r_c_* to be 1.12*σ* to model excluded volume potential. The pairwise electrostatic interaction potential between the *i^th^* bead possessing a charge of *q_i_* and the *j^th^* bead with a charge of *q_j_* separated by a distance *r* is given by the screened Coulombic interaction

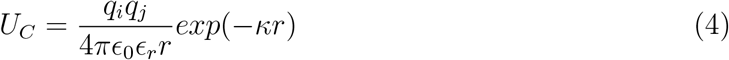

where *ϵ*_0_ is the permittivity of the vacuum, *ϵ_r_* is the dielectric constant of the medium taken to be 60 based on 1.5 M KCl aqueous solution^36^. The inverse Debye length 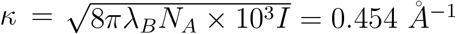, where *I* is the ionic strength in molar, *N_A_* is the Avogadro number and λ_*B*_ is the Bjerrum length given as, λ_*B*_ = *e*^2^/(4*πϵ*_0_*ϵ_r_k_B_T*), *e* being the electronic charge.

The velocities and positions of the protein beads are updated by the rigid body algorithm adopted from Miller *et al*^37^ in LAMMPS^35,38^. The position of the beads belonging to the nanopore are kept fixed. All variables in our simulations are expressed in dimensionless LJ unit, fully consistent with LAMMPS.

### A. Coarse-graining scheme

#### Protein

CG model was built from all atomistic model reported in the PDB (ID: 1UBQ)^30^. The C-α position for each residue was taken as the CG residue center and every residue was assigned a diameter of 3.8 Å^39^. ASP and GLU were assigned −0.2e charge and LYS and ARG were assigned +0.2e charge. The effective charge fraction on the protein is about 0.2 due to the counterion cloud at high salt^27^. At around pH=7, the histidine is neutral^31^, hence it is assigned zero charge.

#### Nanopore

The nanopore parameters are adopted from the experiment^26^. Nanopore was made up of 3.8 A diameter CG beads such that the distance from the center of the pore axis to the bead is 1.99 nm (Pore radius + radius of the CG bead), resulting in an available pore diameter of 3.6 nm for translocation. The length of the pore was originally taken to be 8 nm, but the number of capture events were extremely low for statistical analysis, for few cases. We redefined the problem such that we don’t have to worry about the low capture events. We modeled a nanopore of diameter 3.6 nm and a length of 80 nm (10 times longer than the original nanopore) and started with an initial condition of protein being placed at the center of the nanopore. The dynamics of the protein was then monitored to evaluate the effect of various factors only on the translocation events and not the capture events.

#### Characterization

The results were characterized in three different ways: (a) First Passage Time (*τ*): First time the protein’s center of mass crosses a particular distance starting from an original position at the center of the axis of the pore. (b) z-Mean Square Displacement (MSD): Squared displacement of center of mass from its original position along the nanopore axis. (c) Angular orientation: The angle between the principal axes of Ubiquitin (vector from 76^*th*^ residue to 20^*th*^ residue of PDB) and the nanopore axis. 600 independent simulations were conducted and histogram analysis was done to obtain mean first passage time 〈*τ*〉, and ensemble averaging was done for the other two parameters.

## III. RESULTS AND DISCUSSION

### A. All Atomistic Simulation

The RMSD of the protein backbone was determined as a function of time for two cases: (a) without electric field (b) with applied electric field of 125 mV/ 8 nm applied in any particular orthogonal direction. The Ubiquitin has a radius of gyration (*R_g_*) of about 1.2 nm and the RMSD was determined to be within 2 Å (< 10 % of the diameter of the protein) for all the cases (observation time = 2 ns) (Fig. 1). Since the protein retained its structural integrity as expected for a globular protein, therefore the protein can be approximated as a rigid unit for coarse graining purpose. We are assuming that on an average the instantaneous fluctuations in the conformations of the protein can be neglected for the large time scale simulations. Such approximations may not be appropriate for intrinsic disordered proteins, where the RMSD of the backbone is expected to fluctuate a lot. As the *R_g_*(=1.2 nm) of the protein is smaller than the radius of the nanopore (=1.8 nm), therefore the protein need not deform in order to move inside the nanopore. The protein is elongated (maximum distance between residues is 3.7 nm), hence it cannot enter the nanopore in any orientations. Later in CG simulations, we start with the Ubiquitin principal axis either pointing towards the positive z-direction or the negative z-direction and let it freely evolve into some orientation by keeping it’s center of mass fixed at the initial position, before starting the simulation. In CG simulations we also investigated the effect of electro-osmosis on the dynamics of Ubiquitin. One may ask, what is the impact of the shear on the conformation of the Ubiquitin? At our operational shear rate, does Ubiquitin deform or unravel? According to the all atomistic simulation studies by Walinda et al^40^ on Ubiquitin, upto a shear rate of 1.4× 10^9^ s^-1^, the protein do not show internal structural fluctuations in 20 ns simulation time. In our study, the maximum shear rate is two orders of magnitude smaller, hence our assumption of rigid body model for Ubiquitin seems reasonable in these conditions. Moreover, there is strong experimental evidence that small proteins can travel through SiN_*x*_ nanopore in folded conformation^41^.

**FIG. 1:**
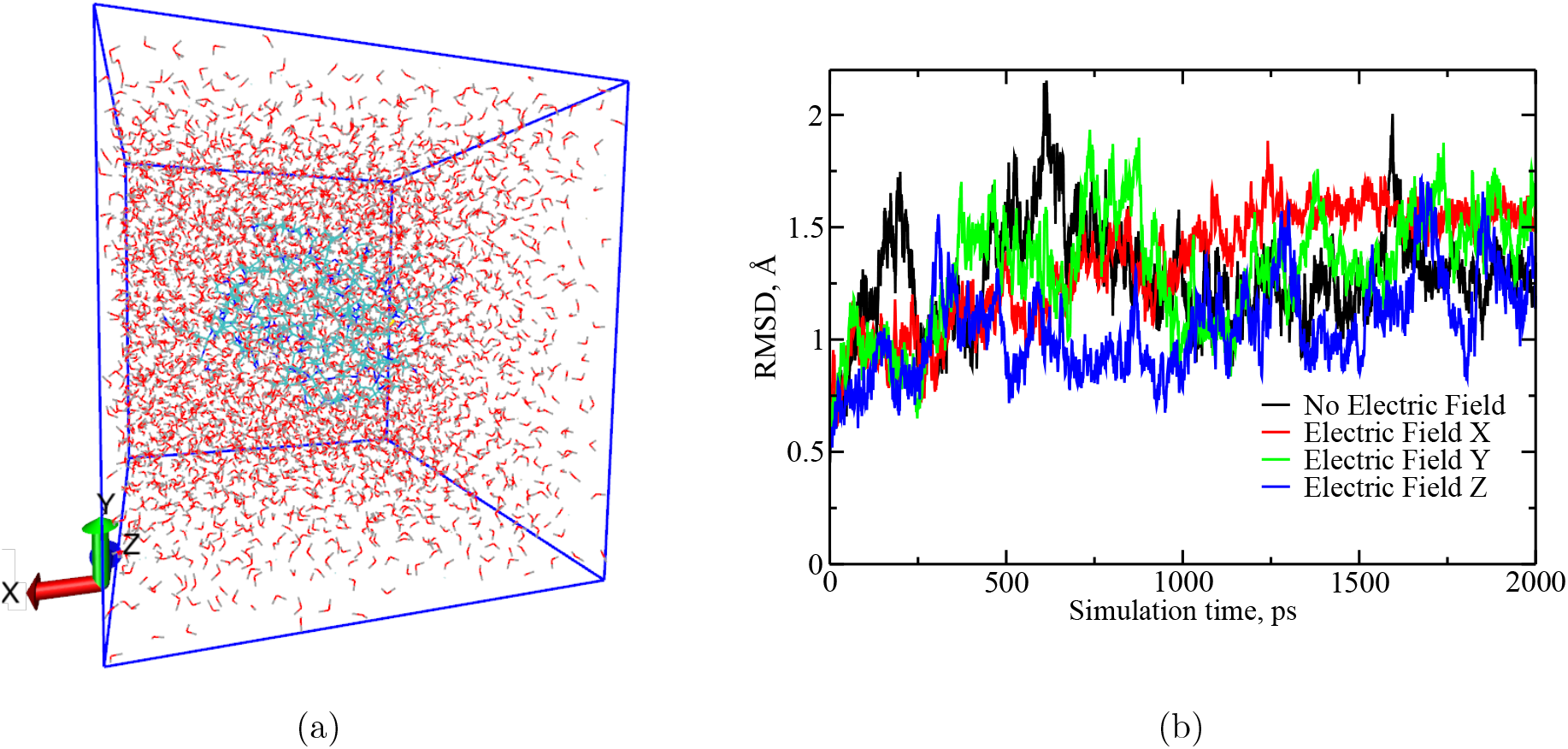
All atomistic MD simulation results demonstrating the stability of Ubiquitin molecular structure in bulk saline solution, irrespective of the presence of an external electric field (|*E*| =0.0015625 V/Å). (a) Snapshot of a Ubiquitin molecule immersed in 1.5 M KCl aqueous solution. The blue stick model in the center of the simulation box is a Ubiquitin molecule which is surrounded by water and dissociated KCl ions. (b) RMSD of the protein backbone as a function of time shows that the fluctuations are within 2Å for all the cases which is less than 10% of the diameter of the protein (= 2x*R_g_* = 24 *Å*), in a simulation time of 2000 ps. Different cases are: no electric field and constant electric field in any orthogonal direction.

### B. Effect of pH on the Protein Charge

The protein is made up of 76 residues, out of which 11 are positive and 11 are negative and one Histidine. Since the pH of the medium is between 7.0-7.2, therefore the charge of Histidine is expected to be zero^31^. The net charge of the protein is zero and for the given experimental conditions (pH =7.0 and pH=7.2), the charge of the protein is not expected to change. Therefore, the observed dramatic change in translocation time due to small change in pH cannot be attributed to modification in the protein charge^26^.

### C. Effect of pH on the Nanopore Surface

In order to determine the effect of pH on the surface charge of SiN_*x*_, experimental literature, Ref.^42^ was followed. According to Ref.^42^, the SiN_*x*_ surface potential/surface charge are 3.6 mV/0.14 *μCcm*^-2^ and −7.2 mV/ −0.11 *μCcm*^-2^ corresponding to pH values of 7.0 and 7.2, respectively, at 0.01 M NaCl. At 1.5 M KCl, we expect the magnitude of the surface potential and surface charge to be reduced, based on their trend for two different salt concentrations: 0.001 M and 0.01 M NaCl. Is it possible to observe such a dramatic change in translocation time distribution with such small values of surface potential and surface charge? Since, we do not know the exact values of these parameters, hence we systematically vary the surface potential between 5 mV to −5 mV.

When the simulation was conducted with the original nanopore length (8 nm), by initiating Ubiquitin near the nanopore mouth, the number of capture events was very low for certain cases. Since Ubiquitin consists of equal number of positive and negative residues, therefore driving it into the nanopore with just electrophoretic force can be challenging. Unless driven by electro-osmosis, the total number of successful capture events were extremely low for further statistical analysis. Since the goal of our study was to study the effects of electrostatic parameters on the dynamics of translocation (not capture), we conducted our simulation in a very long nanopore (80 nm), with the Ubiquitin initiated at the center of the nanopore. The dynamics of the Ubiquitin translocation was investigated for two different sets of forces: (a) protein-pore electrostatic interactions (b) electro-osmosis effects from the charged nanopore.

**FIG. 2:**
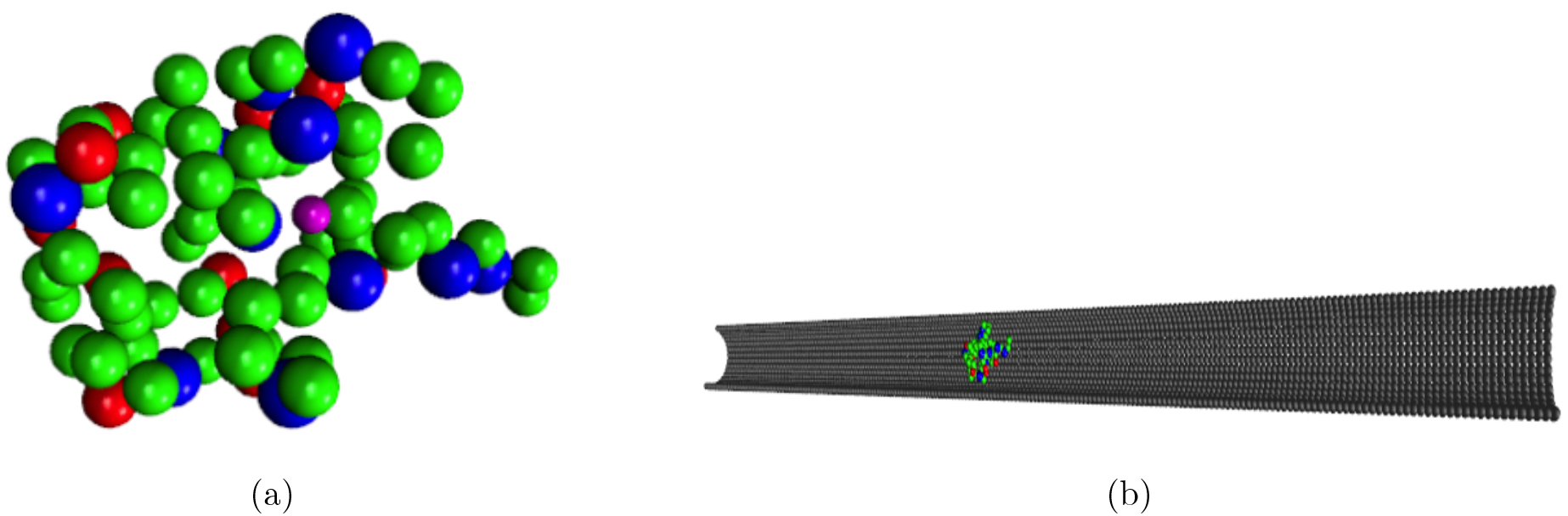
Coarse grain (CG) model of Ubiquitin and the nanopore. (a) Residue based CG model of Ubiquitin derived from it’s all atomistic coordinates (PDB ID: 1UBQ^30^), protein is modeled as a rigid-body. Different residues are color coded based on their charge, neutral: green; positive: blue; negative: red, the lone histidine residue is represented in magenta color. In total there are 76 residues, including 11 positive residues and 11 negative residues, Histidine being neutral at pH = 7.0±0.2^31^. (b)Initial configuration of Ubiquitin in the simulation box, centered in a CG SiN_*x*_ nanopore of 3.6 nm diameter and 80 nm length. The model nanopore diameter was same as the experiment but it’s length was ten times larger^26^. Such a model setup allows understanding of the role of different driving forces on the dynamics of translocation of Ubiquitin by overcoming extremely low capture rate corresponding to certain experimental pH conditions.

#### 1. Protein-pore Electrostatic Interactions

We only consider electrostatic interactions, but the non-specific interactions between the protein molecule and the nanopore are omnipresent. We assume that the small change in pH from 7.0-7.2, modulates only the electrostatic environment inside the nanopore. We considered the simplified case of short-range excluded volume interaction between the residues and nanopore beads and how the additional electrostatic interactions impact the Ubiquitin dynamics. If there is any impact at such simplified conditions then its worth considering more complex short-range interactions. We determined the charge on individual nanopore CG beads (*q_n_*) from Ref.^42^ to be about −0.001e at 7.2 pH. We investigated how the screened coulombic interactions between the charged residues and nanopore affects the first passage time of Ubiquitin COM along the nanopore axis for three different *q_n_* = 0, −0.001e, and −0.01e. Since the Debye length (2.2 Å) is smaller than the CG diameter of a residue (3.8 Å), hence we do not expect any impact of electrostatic interactions. As shown in Fig. 3(a), we observed that the histograms for the three *q_n_* values are very similar. In fact for all the *z* values along the nanopore, the histograms were quite similar, which is evident from the plot of mean first passage time (*τ*) versus *z* (Fig. 3(b)). With increase in *z, τ* increased linearly and symmetrically across the nanopore plane, and *τ* corresponding to *z* = 8nm is about 4 *μs*. This demonstrates that the electrostatic interaction is not responsible for such a drastic change in the average translocation time in response to a small change in pH in the experiment. Next, we investigated the role of electro-osmosis on the mean first passage time.

**FIG. 3:**
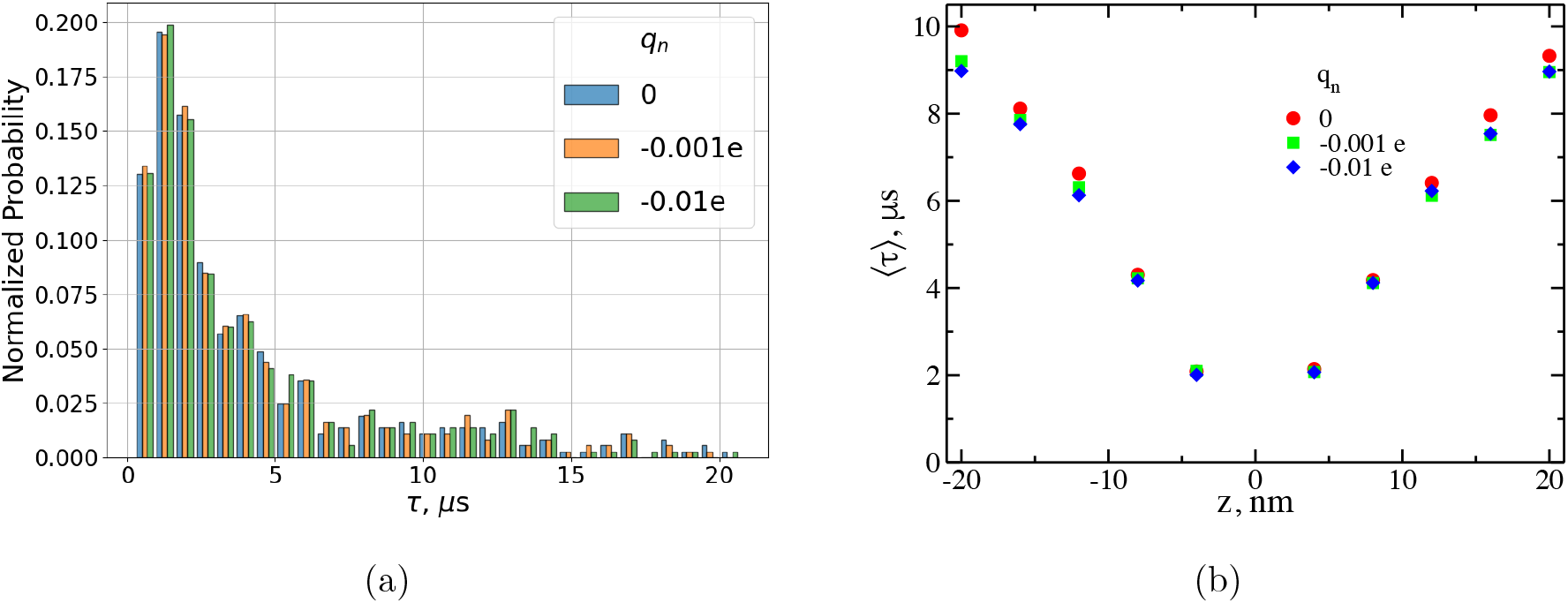
Effects of electrostatic interaction between protein-nanopore on the dynamics of electrophoretically driven Ubiquitin translocation. The charge per CG nanopore bead (*q_n_*) is −0.001e at pH=7.2, we also studied the effect at higher surface charge. (a) A histogram for the first passage time of Ubiquitin at +8 nm, initiated at the center of the nanopore, shows insignificant effect of *q_n_.* Other histograms at +4 nm intervals also showed qualitatively similar behavior. (b) The mean first passage time obtained at regular intervals along the nanopore axis is almost symmetric across the nanopore central plane for all the cases and is insensitive to the charge for any given *z* of the nanopore at such high salt concentration of 1.5 M KCl, as expected.

#### 2. Electro-osmosis Effects

Using the analytical expression derived in Ref.^33^, the fluid velocity profile inside (vf) the nanopore is obtained as shown in Eq. 5, where *ϕ*_0_ is the surface potential, E is the applied electric field, e is the dielectric of the medium, *η* is the viscosity of the medium, *I*_0_ is the zeroth order Bessel function of the first kind, *r* is the radial distance from the pore axis and *a* is the radius of the nanopore. Since the pore is very long, therefore the velocity profile is assumed to be independent of the position of the axis (fully developed flow assumed throughout the length of the nanopore), and is only a function of the radial distance from the nanopore axis. The fluid velocity obtained for the *i^th^* bead located at *r* along the nanopore axis is multiplied by a factor of *m_i_/τ_damp_* = 3.58 × 10^-12^ kg/s to convert it into electro-osmotic force experienced by the residue. This force is then converted to LJ units (Table I) and polynomial fitted (11^*th*^ order), the polynomial expression is used as an input parameter for the langevin equation (Eq. 1).

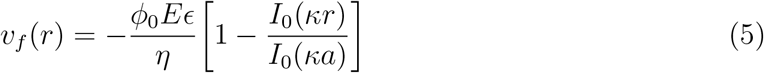

We assumed that the velocity profile of the fluid is not influenced by the presence of Ubiquitin. The fluid velocity *v_f_* is plotted as a function of r for different *ϕ*_0_ as shown in Fig. 4, which is parabolic and its direction is influenced by the magnitude of *ϕ*_0_. For positive values of *ϕ*_0_, *v_f_* is in the negative z-direction, that explains the low capture rate reported in Ref.^26^ at lower pH. At lower pH, the SiN_*x*_ nanopore is expected be more positive^42^. One may ask the applicability of the continuum approximation in such a narrow nanopore. According to Ref.^43^, the ratio of the nanochannel height to approximately the diameter of molecules filling the nanochannel determines whether or not continuum approximations is applicable. By comparing all atomistic molecular dynamics simulations results with the analytical calculations from Navier-Stokes equation, they demonstrated that if this ratio is above 100 then the continuum approximation is valid, below 10 it is not valid, in between the fit between theory and simulation results are not great. Since we are using a nanopore of diameter 3.6 nm, considering diameter of water molecule of 0.27 nm, this ratio is about 13.3. Hence, for our case the continuum approximation is not great, yet not invalid. It should be noted that our velocity is three orders of magnitude smaller than their system, hence the non-linear effects occurring at higher velocities may not be significant in our case. The continuum approximation may be alright for this system, it needs to be tested though.

**FIG. 4:**
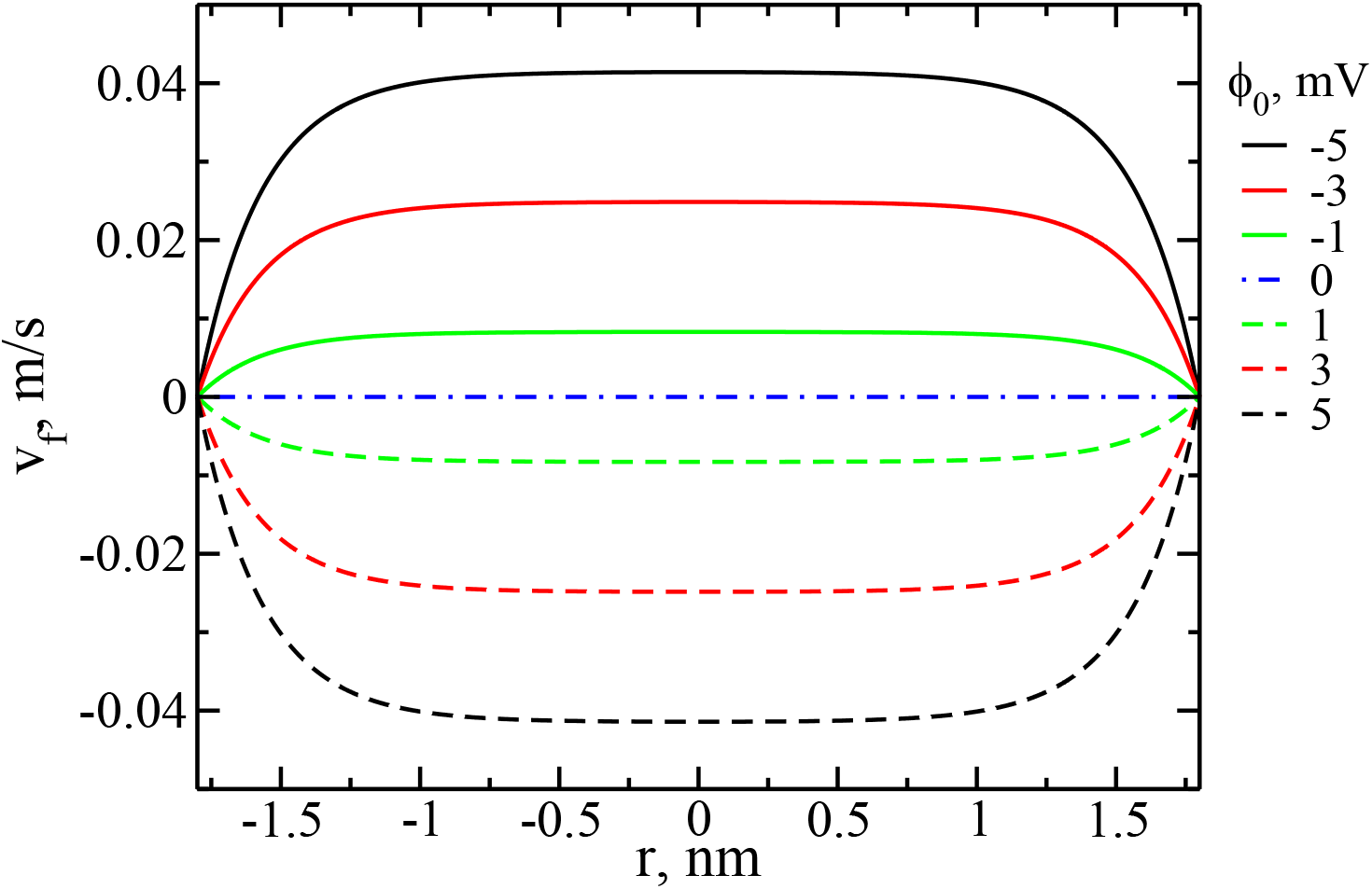
*v_f_* as a function of r is obtained using Eq. 5 for different nanopore surface potentials. The velocity is multiplied by a factor *m_i_/τ_damp_* (in real units) and then converted to LJ force units (Table. I) followed by fitting with a sixth order polynomial function. The polynomial function is used as an input parameter to model electrophoretic force in the langevin equation (Eq. 1).

Next, we investigated the effect of electro-osmosis on the dynamics of Ubiquitin. As shown in Fig. 5(a), there is a drastic effect of electro-osmosis on the first passage time histogram of Ubiquitin at +8 nm along the nanopore axis. The first passage time which spanned till 20 *μs* for *ϕ*_0_ = 0 mV, spans till 1 *μs* for *ϕ*_0_ = −3 mV. The histogram was symmetric on the other side of the nanopore corresponding to positive values of *ϕ*_0_ (not shown). Similar qualitative behavior was observed all along the nanopore as evident from the *τ* versus *z* in Fig. 5(b). *τ* for positive *ϕ*_0_ are not plotted in the positive z-direction, and vice versa, due to insufficient statistics. *τ* versus *z* shows linear behavior for all the cases of *ϕ*_0_. Although we cannot make a direct comparison with the experimental results reported in Ref.^26^ but the (*τ*) values corresponding to *z* = +8 nm are of the the same order of magnitude (~ *μs*) as the dwell time reported in Ref.^26^ for a 8 nm long nanopore, and it decreases with increase in the *ϕ*_0_ value. We do not have enough data to do statistical analysis for positive *ϕ*_0_ values corresponding to *z* =8 nm, but we expect 〈*τ*〉 to increase further with increase in *ϕ*_0_ for longer simulations on a larger sample.

**FIG. 5:**
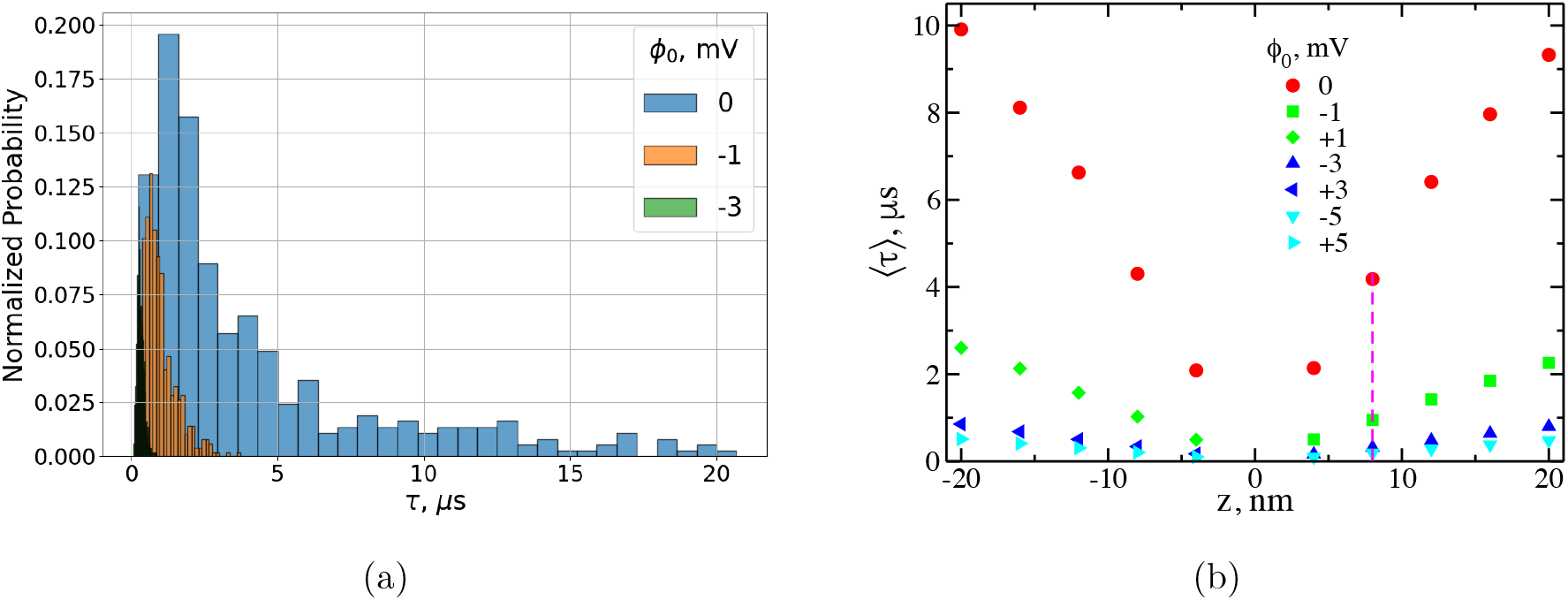
Effect of electro-osmosis on the dynamics of electrophoretically driven Ubiquitin. (a) Histograms for the first passage time of Ubiquitin at +8 nm for three different nanopore surface potentials showing significant decrease in first passage time with decrease in surface potential. For positive surface potentials, there were insignificant number of events corresponding to positive z values, hence not presented. Other histograms at an interval of 4 nm along the nanopore axis are qualitatively similar. (b) The mean first passage time (*τ*) obtained at regular intervals along the pore axis is significantly affected by the surface potential (*ϕ*_0_); higher the magnitude of *ϕ*_0_, lower the *τ*. Negative surface potential biases the protein motion in the positive z-direction, and vice-versa.

We also investigated the effect of *ϕ*_0_ on the z-component of the mean square displacement (z-MSD) as well as the orientation of the protein with respect to the nanopore axis. For the case of zero surface potential, the protein prefers to stay within the nanopore, as evident from a fluctuating z-MSD profile near 100 nm^2^ as shown in Fig. 6(a). For the other cases, z-MSD showed qualitatively different behavior, it increased as a function of time. The slopes of z-MSD increased as the magnitude of *ϕ*_0_ increased. The z-MSD for the positive *ϕ*_0_ values showed very similar trajectories as the corresponding negative *ϕ*_0_ (not shown). Such qualitative trend in z-MSD is expected, based on their first passage time results. Besides translational dynamics, we also investigated the rotational dynamics of Ubiquitin while undergoing translocation through the nanopore. As shown in Fig. 6(b), the orientation of the principal axes (vector from 76^*th*^ residue to 20^*th*^ residue) is independent of *ϕ*_0_ and is dependent only on it’s original orientation at initial time. Although Ubiquitin wiggles a lot (as evident from the thick average value of the orientation) but it cannot flip to the opposite direction due to it’s elongated shape. On an average it prefers an angle of about 34^*o*^ or 150^*o*^ with the nanopore axis depending on it’s initial orientation, probably due to the distribution of charged residues along the protein. We did not observe any effect of the initial orientation on it’s translational dynamics.

**FIG. 6:**
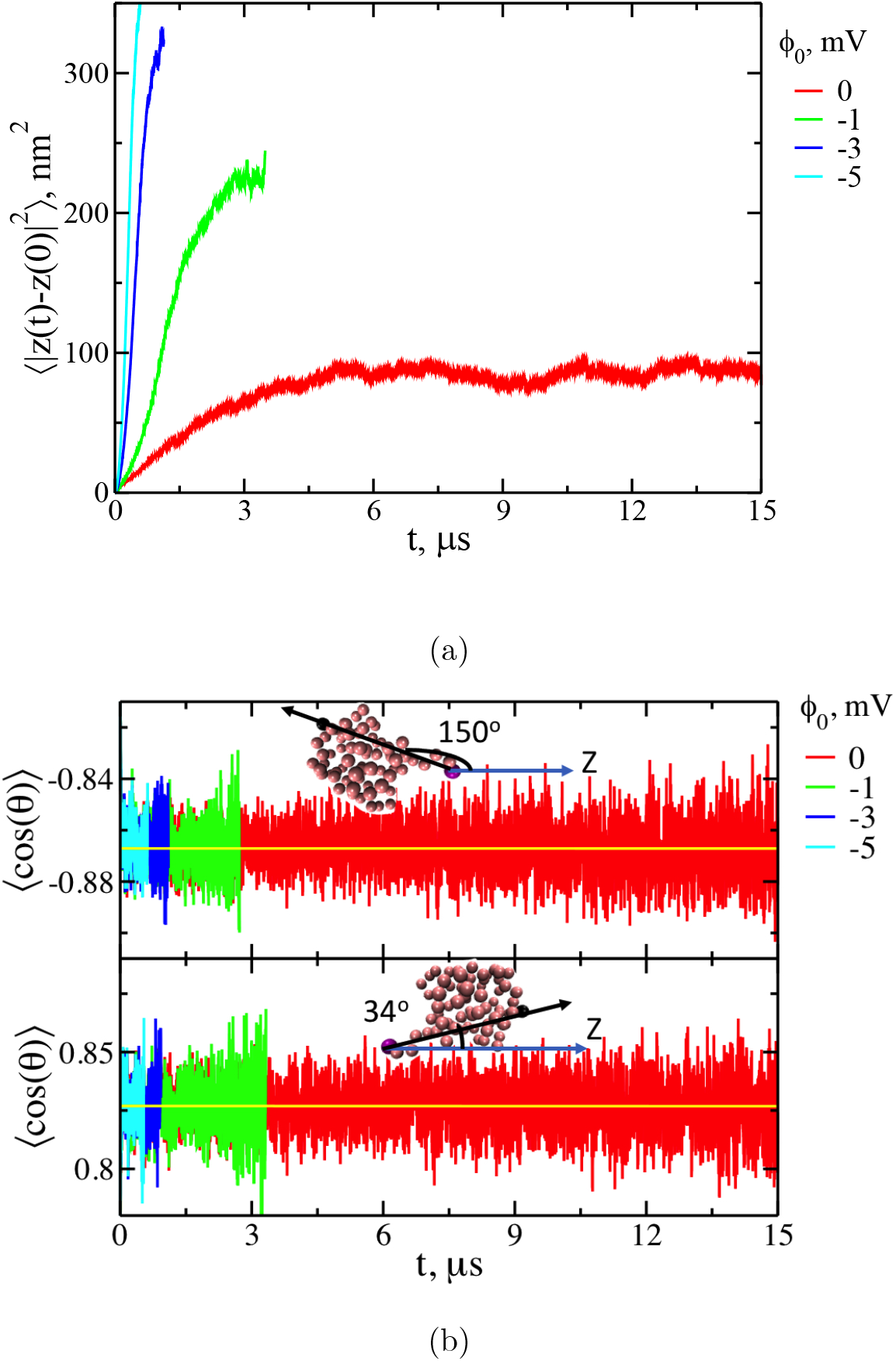
(a) Average of z-component of the mean square displacement (z-MSD) as a function of time showing significant impact of *ϕ*_0_. For *ϕ*_0_=0, the z-MSD steadied around 100 nm^2^, whereas for other *ϕ*_0_, the z-MSD increased drastically as a function of time, showing that drift motion is induced even with slightest increase in surface potential. The z-MSD for positive *ϕ*_0_ are very close to the negative *ϕ*_0_ with the same magnitude (not shown). (b) The orientation of the principal axis of moment (vector from 76^*th*^ residue to 20^*th*^ residue) with respect to z-axis showed preferential orientation along nanopore due to elongated shape of Ubiquitin. The orientation angle is different based on the entry direction of the protein. It does not have much room to flip from its original orientation. small fluctuations in the protein structure can also be included to model a more realistic system without sacrificing too much of computational efficiency.

## IV. CONCLUSIONS

We have developed a framework for determining the large timescale (~ 10 *μs*) dynamics of globular proteins in nanopores. The globular protein can be assumed to be a rigid body, if the protein is smaller along two of its principal axis than the nanopore diameter (so that it fits without getting deformed) and the external force is weak enough such that the conformational fluctuations of the protein can be neglected (high shear can unravel the protein). For cylindrical nanopores of sufficiently large diameter, the electro-osmotic effects can be accounted for by analytical expressions from continuum model. Use of analytical expression for modeling the fluid velocity simplifies the calculations significantly, making it feasible to do simulations of the order of tens of *μs*. With such framework, many different globular proteins can be studied. Investigation can be done on the role of charge distribution and overall charge of the protein on its dynamics. Moreover, the short-range interactions between residue-nanopore pairs can be easily included, if these parameters are known. Lastly, small uctuations in the protein structure can also be included to model a more realistic system without sacrificing too much of computational efficiency.

## V. ACKNOWLEDGEMENT

Acknowledgement is made to the National Institutes of Health (Grant No. 5R01HG002776-16), the National Science Foundation (DMR-2015935), and the AFOSR Grant FA9550-20-1-0142 for financial support.

## Notes

### Competing Interest Statement

The authors have declared no competing interest.

